# BICEP: an extension to indels and copy number variants for rare variant prioritisation in pedigree analysis

**DOI:** 10.64898/2026.03.09.710467

**Authors:** Cathal Ormond, Niamh M Ryan, Aiden Corvin, Elizabeth A Heron

## Abstract

**Summary:** BICEP is a Bayesian inference model that evaluates how likely a rare variant is to be causal for a genomic trait in pedigree-based analyses. The original prior model in BICEP was designed for single nucleotide variants only. Here, we have developed an extension of the prior models for more comprehensive genomic analysis to include indels and copy number variants. We benchmark the performance of these new priors and show comparable performance accuracy with the existing single nucleotide variant prior model. For copy number variants we evaluate four different input predictors to the models and recommend the best performing ones as the default.

**Availability and implementation:** the updated prior models have been implemented in the current version of BICEP available from: https://github.com/cathaloruaidh/BICEP.

## 1 Introduction

Co-segregation analysis has traditionally been used to identify loci, genes, or even specific sequence variants that are implicated in the aetiology of genetic traits (Elandt-Johnson 1971; Bailey-Wilson and Wilson 2011). Many different methodologies and software tools exist for quantifying co-segregation in pedigrees (Almasy and Blangero 1998; Abecasis et al. 2002; Thompson et al. 2003; Mohammadi et al. 2009; Belman et al. 2020; Carrizosa et al. 2024). One limitation shared by these approaches is that they do not consider prior evidence for causality of the variant, such as predicted functional effect on a gene, deleteriousness scores, predicted impact on protein structure, or sequence conservation across species. Variants are usually prioritised by filtering such measures independently of co-segregation. To address this, we developed BICEP, a Bayesian inference model that prioritises candidate causal rare variants by combining co-segregation information with prior evidence for causality (Ormond et al. 2024b). However, these prior models were developed for single nucleotide variants (SNVs) only and were not applicable to other variant types. While SNVs are often the primary focus of analyses using next-generation sequencing data, other variant types such as indels and copy number variants (CNVs) play a role in the aetiology of many complex traits (Beckmann et al. 2007; Auwerx et al. 2022; Hujoel et al. 2022; Wainschtein et al. 2025). Such variants are sometimes ignored in genomic analyses due to challenges in their detection and interpretation (Coughlin et al. 2012), although this limits the amount of publicly available information on contributory risk variation for such traits.

Here we provide an extension of the BICEP prior model for indels and CNVs. As with missense variants and non-missense SNVs in our previous publication, we generated separate prior models for indels, deletions and duplications, since they represent sufficiently different variant types. We benchmark the new prior models with the existing SNV models, showing comparable performance. These updated models have been integrated into the BICEP software and are now available by default.

## 2 Methods

### 2.1 SNV Regression data

In the original version of BICEP, SNV priors were generated using high quality pathogenic and benign variants from the ClinVar database (Landrum et al. 2018), downloaded on 26/11/2023. Here, we have updated the ClinVar downloaded data and evaluated the performance of the SNV prior models in comparison to the indels and CNVs using this updated version of the database, downloaded 08/12/2025. To compare the performance across genome builds, we obtained data on GRCh37 and GRCh38. These datasets were processed identically unless otherwise specified. Similar to the SNV prior odds, all prior odds for indels and CNVs are given on the log10 scale, referred to as a logPriorOC.

### 2.2 Regression data for extension to indels

To generate a prior for indels, we adapted the previous BICEP framework for SNVs. Indels in the ClinVar regression data were allowed to have support of “single submitter” as well as “multiple submitters” to increase the size of the dataset. Additionally, instead of using the predicted functional consequence from VEP as an input to the regression model (McLaren et al. 2016), we used the more broadly defined IMPACT annotation, which groups the consequences into four levels: HIGH, MEDIUM, LOW, and MODIFIER. These two modifications were to ensure that each category had a sufficient minimum number of data points in the regression data. Allele frequencies from gnomAD (Chen et al. 2024) were included as a predictor in the model by default, using v4.1 for data on GRCh38 and v2.1.1 for data on GRCh37.

### 2.3 Regression data for extension to CNVs

For CNVs, clinical structural variants were obtained from the dbVar database (Lappalainen et al. 2013), study ID nstd102, (dated 13/08/2025). As with the SNV and indel model, data were downloaded for both genome builds. These data were used to train a logistic regression model using a pre-specified set of predictors (see below). Like the SNV and indel models, using the regression coefficients, a prior odds for CNV causality (i.e., pathogenicity) can be generated. From the CNV regression data, we subset to deletions and duplications whose length was between 50bp and 30Mbp, consistent with typical CNV calling from whole genome sequencing data (Ormond et al. 2024a). Variants were retained if their clinical significance was “Pathogenic” and/or “Likely Pathogenic”, or “Benign” and/or “Likely Benign”. Where multiple records had the same coordinates, one was retained at random if the clinical significance was the same for all records. Where there was a conflict in the clinical significance, these records were removed. We also removed CNVs that had no overlap with the MANE v1.4 transcript of a protein-coding gene (Morales et al. 2022). The maximum allele frequencies across all genomic ancestry clusters (i.e. PopMax) from gnomAD were annotated using SFAVotate (Nicholas et al. 2022) using v4.1 for data on GRCh38 and v2.1.1 for data on GRCh37, first removing gnomAD variants without a “PASS” filter. The largest allele frequency across all CNVs of matching type (i.e., deletion or duplication) with a 50% reciprocal overlap was selected. However, any pathogenic CNV with a reported allele frequency above 1% was removed from the regression data, as such variants are typically considered to be benign (Riggs et al. 2020).

### 2.4 Predictors

For the indel model, we only considered the default predictors of allele frequency and variant functional consequence. We did not include additional deleteriousness predictors, as many such scores available for indels are trained on ClinVar data (e.g., CAPICE (Li et al. 2020), INDELpred (Wei et al. 2024)), which would result in a circularity bias (Grimm et al. 2015).

For each CNV in the regression data, we generated CADD-SV deleteriousness scores (Kleinert and Kircher 2022). Since CADD-SV can only be run on data generated on build GRCh38, we converted CNVs to GRCh37 using liftOver (Haeussler et al. 2019). Additionally, we aggregated gene-based loeuf constraint scores from gnomAD v4.1 (Karczewski et al. 2020; Chen et al. 2024) for each CNV. For all genes whose MANE transcript was completely contained within the CNV region, we calculated the sum of the reciprocal of the loeuf score (Brownstein et al. 2022), since lower scores indicate intolerance to loss-of-function. The above predictors were annotated using bcftools (Danecek et al. 2021). We examined four models for deletions and duplications separately: no predictors (called the “NULL” model), CADD-SV scores, the loeuf measure, and both CADD-SV and the loeuf measure.

### 2.5 Model benchmarking

We split the regression data into training (80%) and testing (20%) subsets and generated binary classification metrics for the hold-out testing set. A variant was predicted to be “pathogenic” if it had logPriorOC > 0, and it was predicted to be “benign” otherwise. As in the original BICEP publication (Ormond et al. 2024b), we calculated sensitivity, specificity, positive predictive value (PPV), negative predictive value (NPV), and the Matthews correlation coefficient (MCC). We focused on the PPV as our primary benchmark metric, as it represents the proportion of variants with logPriorOC > 0 that are truly pathogenic, which is typically the focus of a BICEP analysis. Analogously, the NPV represents the proportion of variants with logPriorOC < 0 that are truly benign.

### 2.6 Availability

The new prior models and regression data have been integrated into the current implementation of BICEP (https://github.com/cathaloruaidh/BICEP). The default analysis assumes the input is composed of SNVs and/or indels. Users have the option of specifying that the input data are CNVs, and so the appropriate prior model should be used. The same input data formats as described previously (Ormond et al. 2024b) can be used for the new models.

## 3 Results

The performance metrics for the new models on GRCh38 data are shown in (short variants, deletions and duplications). For comparison, we included the metrics for the two SNV models (missense and non-missense SNV) described previously (Ormond et al. 2024b). Similar metrics for data on build GRCh37 can be found in Supplementary Figure 1. Across all variant types and models evaluated, the confidence intervals for metrics from the training overlapped well with those from the testing data (see Supplementary Figure 2), indicating that overfitting is unlikely. We note that for indels, the PPV was very high, at 99.8%. The NPV for indels was modest (62.3%) compared to the two existing SNVs models (>96.0%). This indicates that some truly pathogenic indels have a negative logPriorOC and so are not well characterised by the model. The wider confidence intervals for the indel model likely reflect the smaller size of the regression data, as well as the imbalance between pathogenic and benign variants (see Table 1).

**Table 1:** The number of pathogenic and benign variants in the GRCh38 ClinVar regression data for each of the five BICEP prior models.

| **Variant type** | **Pathogenic** | **Benign** | **Total** |
| --- | --- | --- | --- |
| Missense variant | 12,539 | 24,672 | 37,211 |
| Non-missense SNV | 19,985 | 98,604 | 118,589 |
| Indel | 3,508 | 393 | 3,901 |
| Deletion | 22,398 | 2,317 | 24,715 |
| Duplication | 4,413 | 3,903 | 8,316 |

**Figure 1.**
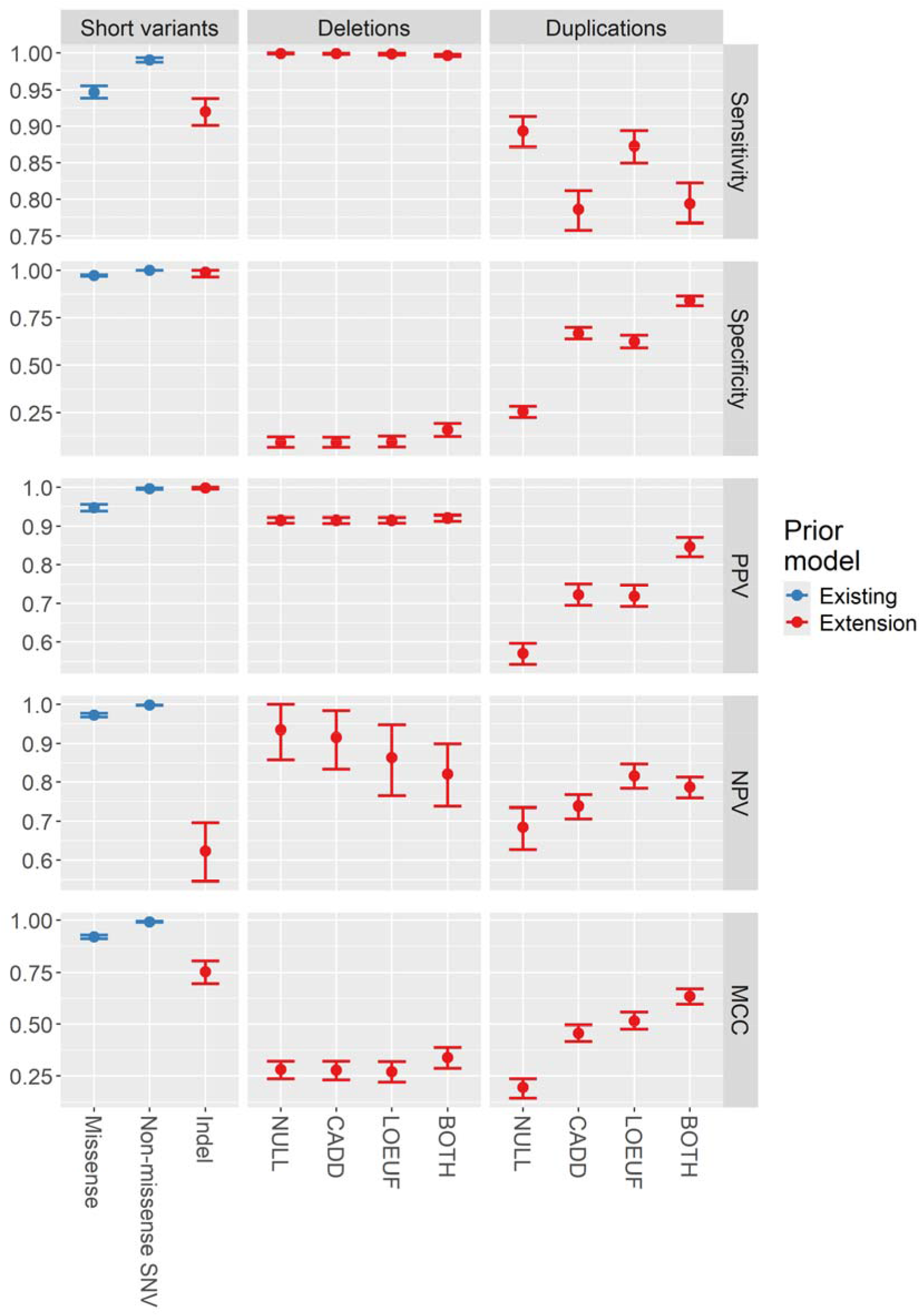
The performance metrics for the various BICEP prior models under evaluation for GRCh38. “NULL” indicates that only allele frequency was used, and “BOTH” indicates that the CADD-SV and loeuf measure were used. The metrics are given for the two existing prior models for SNVs (blue), and for the extensions to indels and CNVs (red). PPV: positive predictive value; NPV: negative predictive value; MCC: Matthews correlation coefficient; SNV: single nucleotide variant.

For deletions, the model using both CADD scores and the loeuf measure had the highest point estimate of the PPV (92.0%), although there was little difference in PPV across the four models evaluated. There was some variability in the NPVs, although we note that each model had a small number of variants with negative logPriorOC values (Supplementary Table 2). Overall, however, the confidence intervals for the four models were largely overlapping. Based on this, we recommend using both CADD-SV scores and the loeuf measure as additional predictors for deletions, (i.e. the BOTH model). For duplications, the model using both predictors had the highest PPV (84.6%), which noticeably outperformed the other three models evaluated. While the NPV for this model was lower than one of the others, it still performed comparably well, and so we recommend using both scores as additional predictors for duplications.

## 4 Discussion

We have described an extension of the BICEP prior model for indels and CNVs. We showed that the new models perform comparably with the existing SNV models. The NPVs for the extended models indicate that some truly pathogenic variants are given a negative logPriorOC score. While more training data as well as better deleteriousness predictors will improve this, the PPV for these models already performs reasonably well. Since a typical BICEP analysis focuses on variants with positive logPriorOC scores, any changes are unlikely to markedly improve the PPV. In the two CNV models, the PPV for the best performing set of predictors was slightly lower than those for the short variants. However, this can be explained in part by the substantially fewer number of deleteriousness predictors available for CNVs compared to SNVs.

As indels and CNVs are of interest in complex diseases it is important that BICEP has the capacity to analyse these variant types. We acknowledge that the performance for these variant types is not as high as for the SNVs, although this is not unexpected given that SNVs have historically been more predominantly studied. We expect the prior models for indels and CNVs to improve as better predictors are developed. Also, as more research groups turn to modern sequencing technologies, the size and quality of the regression data will increase, which will facilitate retraining. Hence it is essential to have the models available at this stage.

## Acknowledgements

This work was supported by National Institutes of Health [5R01MH 124875 to A.C.].

