## Supplementary materials for "BICEP: an extension to indels and copy number variants for rare variant prioritisation in pedigree analysis"

### 1 Supplementary figures

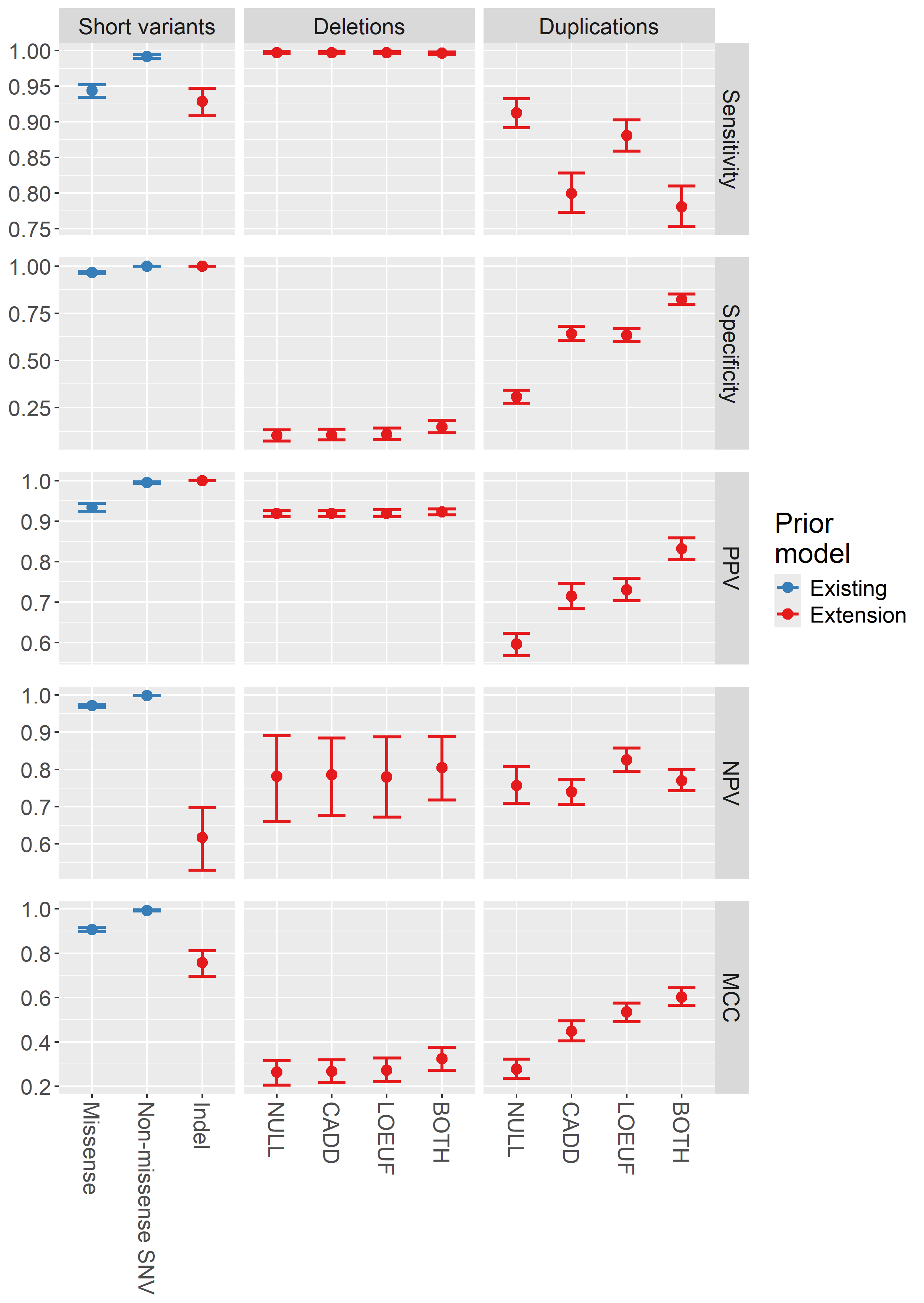

Supplementary Figure 1: The performance metrics for the various BICEP prior models under evaluation for GRCh37. “NULL” indicates that only allele frequency was used, and “BOTH” indicates that the CADD-SV and aggregated loeuf scores were used. The metrics are given for the two existing prior models for SNVs (blue), and the extensions to indels and CNVs (red). PPV: positive predictive value; NPV: negative predictive value; MCC: Matthews correlation coefficient; SNV: single nucleotide variant.

**(A)**

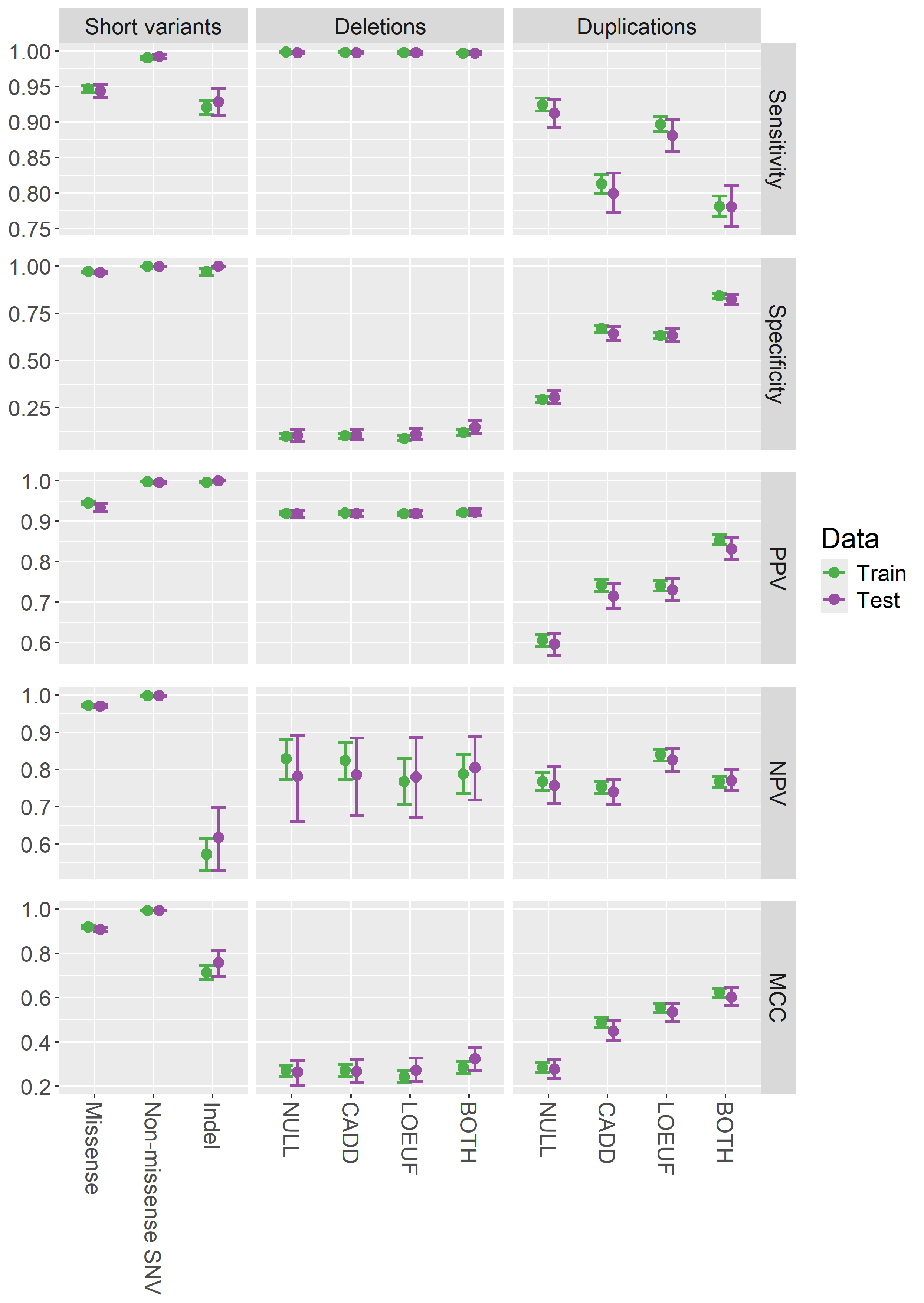

**(B)**
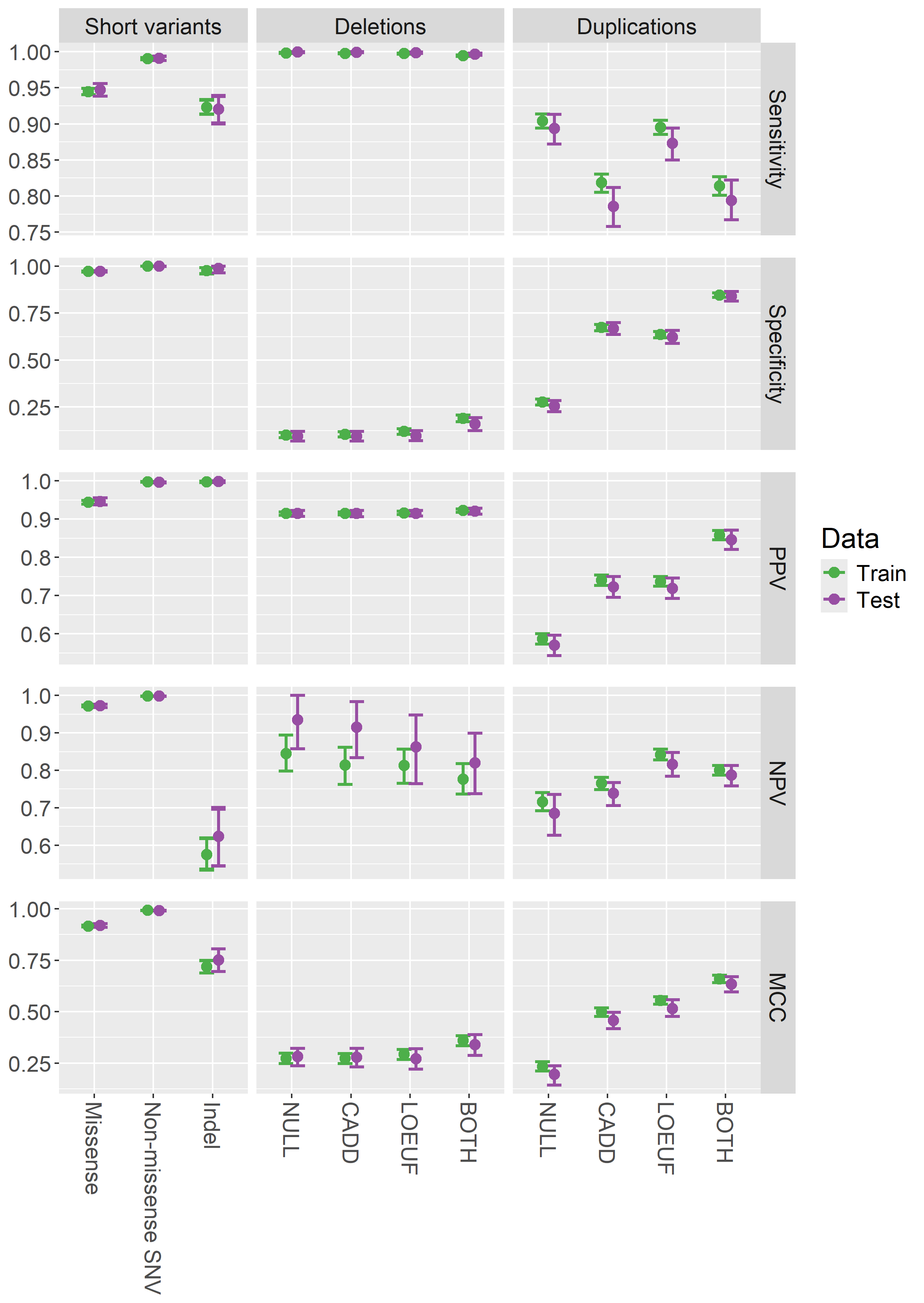

Supplementary Figure 2: The performance metrics for the various BICEP prior models on **(A)** GRCh37 and **(B)** GRCh38. “NULL” indicates that only allele frequency was used, and “BOTH” indicates that the CADD-SV and aggregated loeuf scores were used. The metrics are given for the training data (green) and the hold out test data (purple). PPV: positive predictive value; NPV: negative predictive value; MCC: Matthews correlation coefficient; SNV: single nucleotide variant.

### 2 Supplementary tables

| **Variant type** | **Pathogenic** | **Benign** | **Total** |
| --- | --- | --- | --- |
| Missense variant | 12,266 | 23,814 | 36,080 |
| Non-missense SNV | 19,560 | 96,461 | 116,021 |
| Indel | 3,442 | 380 | 3,822 |
| Deletion | 21,492 | 2,081 | 23,573 |
| Duplication | 4,053 | 3,489 | 7,542 |

Supplementary Table 1: The number of pathogenic and benign variants in the GRCh37 ClinVar regression data for each of the five BICEP prior models.

**(A)**

|  |  | **GRCh37** | | | **GRCh38** | | |
| --- | --- | --- | --- | --- | --- | --- | --- |
|  | **logPriorOC** | **BEN** | **PATH** | **TOTAL** | **BEN** | **PATH** | **TOTAL** |
| **NULL** | ≥0 | 378 | 4,282 | 4,660 | 417 | 4,480 | 4,897 |
|  | <0 | 43 | 12 | 55 | 43 | 3 | 46 |
|  | Total | 421 | 4,294 | 4,715 | 460 | 4,483 | 4,943 |
| **CADD** | ≥0 | 377 | 4,282 | 4,659 | 417 | 4,479 | 4,896 |
|  | <0 | 44 | 12 | 56 | 43 | 4 | 47 |
|  | Total | 421 | 4,294 | 4,715 | 460 | 4,483 | 4,943 |
| **LOEUF** | ≥0 | 375 | 4,281 | 4,656 | 416 | 4,476 | 4,892 |
|  | <0 | 46 | 13 | 59 | 44 | 7 | 51 |
|  | Total | 421 | 4,294 | 4,715 | 460 | 4,483 | 4,943 |
| **BOTH** | ≥0 | 359 | 4,279 | 4,638 | 387 | 4,467 | 4,854 |
|  | <0 | 62 | 15 | 77 | 73 | 16 | 89 |
|  | Total | 421 | 4,294 | 4,715 | 460 | 4,483 | 4,943 |

**(B)**

|  |  | **GRCh37** | | | **GRCh38** | | |
| --- | --- | --- | --- | --- | --- | --- | --- |
|  | **logPriorOC** | **BEN** | **PATH** | **TOTAL** | **BEN** | **PATH** | **TOTAL** |
| **NULL** | ≥0 | 493 | 728 | 1,221 | 589 | 780 | 1,369 |
|  | <0 | 218 | 70 | 288 | 202 | 93 | 295 |
|  | Total | 711 | 798 | 1,509 | 791 | 873 | 1,664 |
| **CADD** | ≥0 | 255 | 638 | 893 | 263 | 686 | 949 |
|  | <0 | 456 | 160 | 616 | 528 | 187 | 715 |
|  | Total | 711 | 798 | 1,509 | 791 | 873 | 1,664 |
| **LOEUF** | ≥0 | 260 | 703 | 963 | 298 | 762 | 1,060 |
|  | <0 | 451 | 95 | 546 | 493 | 111 | 604 |
|  | Total | 711 | 798 | 1,509 | 791 | 873 | 1,664 |
| **BOTH** | ≥0 | 126 | 623 | 749 | 126 | 693 | 819 |
|  | <0 | 585 | 175 | 760 | 665 | 180 | 845 |
|  | Total | 711 | 798 | 1,509 | 791 | 873 | 1,664 |

Supplementary Table 2: For the four CNV prior models, counts of variants in the test dataset broken down by whether their logPriorOC is negative or non-negative for **(A)** deletions and **(B)** duplications.
